# A fur plucking model to study herpes simplex virus reactivation and recurrent disease

**DOI:** 10.1101/2023.12.06.570385

**Authors:** Drake T. Philip, Nigel M. Goins, Helen M. Lazear

**Affiliations:** Department of Microbiology & Immunology, University of North Carolina at Chapel Hill

## Abstract

Herpes simplex viruses (HSV-1 and HSV-2) most commonly cause ulcerative epithelial lesions (cold sores, genital herpes). Importantly, HSV establishes life-long persistent (latent) infection in sensory neurons. Reactivation from latency produces recurrent epithelial lesions, which constitute the greatest burden of HSV disease in people. The mechanisms that regulate latency and reactivation remain poorly understood, in part due to limitations in the animal models available for studying HSV reactivation. We have developed a simple and tractable model to induce HSV-1 and HSV-2 reactivation from latently infected sensory ganglia. We infected C57BL/6 mice with 1 × 10^6^ FFU of HSV-1 (strain NS) or 500 FFU of HSV-2 (strain 333) on flank skin depilated by manual plucking. 35 days post-infection (dpi) we re-plucked the fur from the infected flank and observed recurrent lesions in the same dermatome as the primary infection. We detected HSV DNA in dermatome skin through 4 days post-re-plucking. We found that shaving the ipsilateral flank or plucking the contralateral flank did not induce recurrent skin lesions, suggesting that fur plucking is a specific stimulus that induces HSV reactivation. Further, we were able to induce multiple rounds of plucking induced recurrent disease, providing a model to investigate the lifelong nature of HSV infection. This new model provides a tractable system for studying pathogenic mechanisms of and therapeutic interventions against HSV reactivation and recurrent disease.

## IMPORTANCE

Herpes simplex viruses (HSV-1 and HSV-2) are highly prevalent and cause lifelong persistent infections. However, our understanding of the mechanisms that govern HSV reactivation and recurrent disease are limited in part due to poor animal models to study recurrent disease, which typically require corneal infection and the methods to induce viral reactivation are laborious or inefficient. To address this, we developed a mouse model in which fur plucking induces HSV reactivation and recurrent disease after skin infection. Our work provides a model for the field to investigate the pathogenic mechanisms of HSV and immune responses during recurrent disease and provides an opportunity to investigate the neurobiology of HSV infection.

## OBSERVATION

Herpes simplex virus type 1 and type 2 (HSV-1, HSV-2) are significant human pathogens infecting over half of the US adult population (1). Following infection at epithelial surfaces (e.g. skin, cornea), HSV causes lifelong persistent infections by establishing latency in sensory neurons. HSV can then reactivate from latency and travel via anterograde axonal transport to the epithelium where it produces recurrent lesions and is transmitted to new individuals (2, 3). While HSV infection most commonly results in orofacial or genital lesions, severe manifestations of HSV infection include encephalitis, keratitis, and neonatal infections (2). The greatest burden of HSV disease results from the ability of HSV to reactivate and cause recurrent disease throughout the lifetime of an infected person but the tools available to study mechanisms of reactivation and recurrent disease are limited, especially with regard to mouse models of recurrent disease. Most animal models used to study HSV reactivation are based on corneal inoculation, where recurrent disease is not straightforward to evaluate (4, 5). Further, HSV does not spontaneously reactivate in mice, and the stimuli used to induce reactivation, such as ultraviolet light (6), hyperthermia (7), stress (8), and hormone treatment (9), are inefficient and have limited tractability. Therefore, we sought to develop a more tractable model to study mechanisms of HSV reactivation and recurrent disease.

To study acute HSV disease, we used the flank infection model (10, 11) optimized to improve operator-to-operator and mouse-to-mouse variability (12). In brief, one day prior to infection, we depilated mice by manual fur plucking, rather than depilating with shaving and Nair cream. In this model, the virus replicates in the skin at the inoculation site, infects innervating sensory neurons, traffics to the neuron cell bodies in the dorsal root ganglia (DRG), then returns to the entire dermatome innervated by that DRG, producing a skin lesion. We measure the area of dermatome skin lesions from photographs and measure viral loads in the skin lesions by qPCR. We previously found that depilation by plucking had no effect on skin lesion area or viral loads compared to depilation by shave/Nair (12). However, based on prior studies that used tape stripping-induced reactivation with an ear pinna infection model (13), we asked whether fur plucking was sufficient to induce HSV reactivation and recurrent disease.

To test whether fur plucking can induce HSV reactivation, we infected C57BL/6 (wild-type) mice with 1 × 10^6^ focus-forming units (FFU) of HSV-1 strain NS, evaluated their acute skin disease 6 days post-infection (dpi), then allowed the infection to resolve, the fur to regrow, and the virus to establish latency. 5 weeks after the infection we re-plucked the ipsilateral flank and evaluated skin lesions through 6 days post-reactivation (R6). While no dermatome lesions were evident immediately upon re-plucking (recurrent day 0, R0), consistent with immune-mediated clearance of the acute lesion and establishment of viral latency, by 2 days post-reactivation (R2) mice exhibited skin lesions in the same dermatome as their acute disease. These recurrent lesions resolved by 6 days post-reactivation (R6) (Fig. 1A-B). To determine whether recurrent lesions corresponded to HSV infection in dermatome skin, we infected wild-type mice with HSV-1, re-plucked 35 dpi, collected dermatome skin from R0 to R6, and measured HSV-1 DNA by qPCR (Fig. 1C-D). While only 1 of 15 mice had detectable viral DNA in the skin at R0, by R2 we detected HSV-1 DNA in the skin of 8 of 16 mice. At R4 the frequency of HSV-1 positive animals was similar (11 of 22), but the maximum viral load of the positive samples had increased (3.27 Log_10_ FFU equivalents at R4 compared to 2.38 at R2). All mice cleared HSV-1 from the skin by R6. The relatively low viral loads observed are consistent with other models of HSV reactivation and with the presence of adaptive immune responses at this stage of infection. These results suggest that fur plucking induced HSV-1 to reactivate in latently-infected DRG and traffic to dermatome skin, where it replicated and produced a lesion.

**Figure 1.**
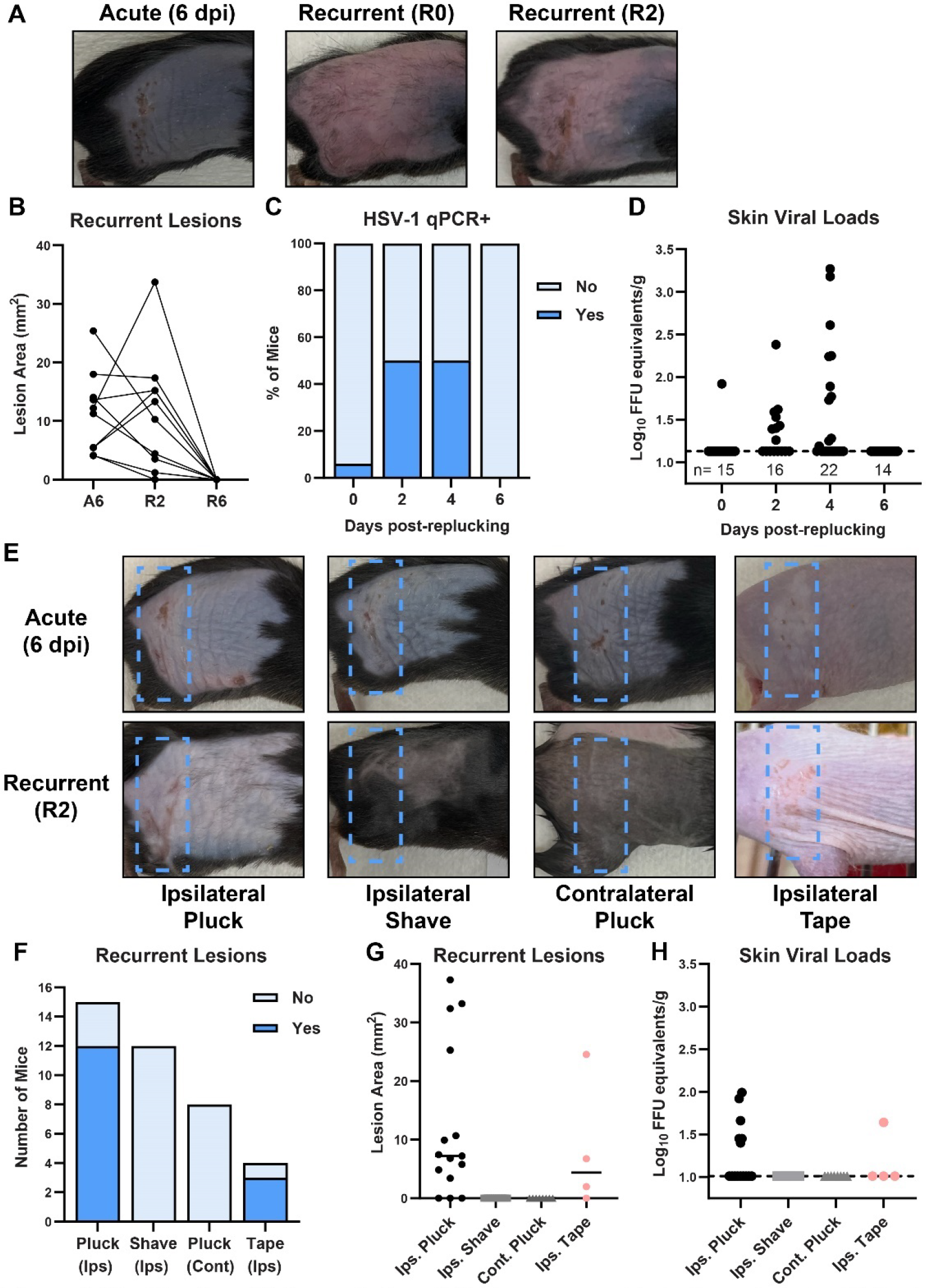
Plucking is sufficient to induce HSV-1 recurrent skin lesions. A-D. C57BL/6 mice were depilated by plucking one day prior to infection then inoculated on the flank skin with 1 × 10^6^ FFU of HSV-1 strain NS. 35 days later, mice were replucked and recurrent skin lesions were evaluated. **A**. Representative photos of the same mouse during acute disease (6 dpi), immediately post-replucking (RO), and 2 days post-repluck ing (R2). **B**. Paired lesion areas of mice during acute infection, 2 days post-replucking, and 6 days post-replucking. **C-D**. Viral loads in dermatome skin post-replucking were measured by qPCR; dotted line indicates limit of detection. **E-H**. C57BL/6 mice were depilated by plucking one day prior to infection then inoculated on the flank skin with 1 × 10^6^ FFU of HSV-1 strain NS. Hairless (SKH-1) mice were inoculated with 5 × 10^6^ FFU. 35 days later dermatome skin lesions on the ipsilateral flank were evaluated after: replucking the ipsilateral flank; shaving the ipsilateral flank; replucking the contralateral flank (plus shaving the ipsilateral flank); or tape stripping the ipsilateral flank. **E**. Representative photos of the same mouse during acute disease (6 dpi) and 2 days post-replucking (R2). **F-G**. Incidence and area of recurrent lesions 2 days after stimulation. **H**. Viral loads in dermatome skin 2 days after stimulation were measured by qPCR.

We next asked whether fur plucking per se induced HSV-1 reactivation, or whether reactivation might be induced by the stress of handling/manipulating the mice or temperature changes due to fur removal. To investigate this, we infected wild-type mice with HSV-1 and allowed the virus to establish latency. 35 dpi we re-depilated the ipsilateral flank by plucking or by shaving with clippers. While 12 of 15 re-plucked mice exhibited recurrent skin lesions and 6 had HSV-1 DNA present in the skin by qPCR, 0 of 12 re-shaved mice exhibited recurrent skin lesions and no HSV-1 was detected in the dermatome skin (Fig. 1 F-H), indicating that shaving was not a sufficient stimulus to induce reactivation. We next tested whether plucking stimulated reactivation by inducing a systemic response or a local one. We infected wild-type mice with HSV-1 and 35 dpi we re-plucked the contralateral flank and shaved the ipsilateral flank to allow recurrent lesions to be observed. From 8 mice re-plucked contralaterally we observed no recurrent dermatome lesions and no HSV-1 DNA was detected in the skin (Fig. 1F-H), indicating that the plucking stimulus had to be at the ipsilateral flank to induce reactivation. To further investigate the ability of hair removal to induce HSV-1 reactivation, we used SKH-1 mice, an immunocompetent mouse line commonly described as “hairless” but which actually have very fine hair (14). We infected SKH-1 mice with 5 × 10^4^ FFU of HSV-1 strain NS. 4 of 9 mice survived the infection (these mice are more susceptible to HSV-1 compared to C57BL/6 mice) and 35 dpi we removed their ipsilateral flank hair by stripping with autoclave tape. We found that 3 of 4 tape-stripped SKH-1 mice developed recurrent lesions by R2 (Fig. 1E-G) and one had detectable HSV-1 DNA in the skin at R2 (Fig. 1H). Altogether, these data support a model where fur plucking stimulates reactivation of latent HSV-1 from sensory neurons innervating that site, resulting in recurrent skin lesions.

A defining feature of HSV disease in humans is its ability to cause multiple recurrent disease episodes throughout the lifetime of an infected individual (2). Therefore, we next asked whether fur plucking could stimulate multiple rounds of reactivation and recurrent disease. We infected wild-type mice with 1 × 10^6^ FFU of HSV-1 strain NS and evaluated acute disease 6 dpi. We then re-plucked the mice 35 dpi, evaluated recurrent skin lesions on R2, then let the mice recover again and regrow their fur. Then, 35 days after re-plucking (RR0), we re-plucked the mice again and evaluated skin lesions. As expected, no skin lesions were evident immediately upon re-plucking (RR0), indicating that the prior recurrent skin lesion had cleared. However, after 2 days (RR2) we found that 9 of 11 mice exhibited recurrent skin lesions in the same dermatome as their acute lesions and their first recurrent lesions (Fig. 2A-B). Altogether, these data demonstrate that fur plucking can cause multiple rounds of HSV-1 reactivation in mice.

**Figure 2.**
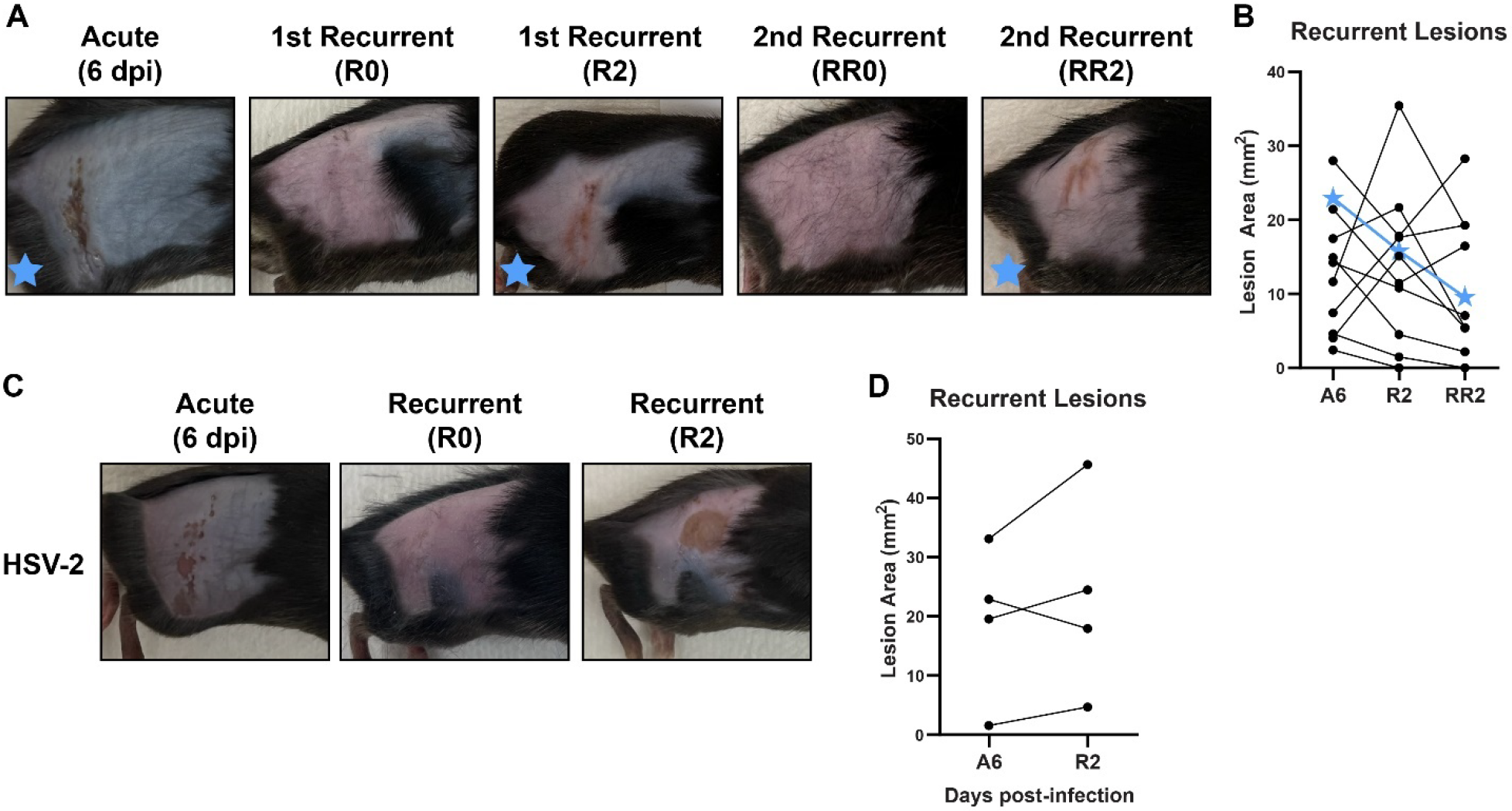
Plucking can induce multiple rounds of reactivation and can reactivate both HSV-1 and HSV-2. A-B. C57BL/6 mice were depilated by plucking one day prior to infection then inoculated on the flank skin with 1 × 10^6^ FFU of HSV-1 strain NS. 35 days later, mice were replucked and recurrent skin lesions were measured. After another 35 days, mice were replucked again and recurrent skin lesions were measured. **A**. Representative photos of the same mouse during acute disease (6 dpi), immediately post-replucking (R0 and RR0), and 2 days post-replucking (R2 and RR1). **B**. Paired lesion areas of mice during acute infection and 2 days post-replucking. Star symbols indicate the mouse shown in (**A**). **C-D**. C57BL/6 mice were depilated by plucking one day prior to infection then inoculated on the flank skin with 5 × 10^2^ FFU of HSV-2 strain 333. 35 days later, mice were replucked and recurrent skin lesions were measured. **C**. Representative photos of the same mouse during acute disease (6 dpi), immediately post-replucking (R0), and 2 days post-replucking (R2). **D**. Paired lesion areas of mice during acute infection and 2 days post-replucking.

In addition to HSV-1 strain NS, we were able to similarly reactivate HSV-1 strain SC16 and strain F (data not shown). We next asked whether this reactivation model was specific to HSV-1 or more broadly applicable to HSV-2 as well. We infected wild-type mice with 500 FFU of HSV-2 strain 333 and evaluated lesions 6 dpi. We used a lower dose of HSV-2 than HSV-1 because at higher doses most mice succumbed to HSV-2 infection, precluding reactivation studies. At a dose of 500 FFU, 8 of 14 mice exhibited dermatome lesions at 6 dpi and 4 of these survived the infection. We allowed these 4 mice to recover from the acute infection and regrow their fur, then re-plucked them 35 dpi. We found 4 of 4 mice developed recurrent lesions after re-plucking (Fig. 2 C-D). Altogether, these data show that fur-plucking can induce reactivation and recurrent skin disease for both HSV-1 and HSV-2.

Overall, we report a new mouse model to study HSV reactivation and recurrent disease, in which plucking fur is sufficient to stimulate HSV reactivation from latently infected DRG. This new model has several advantages compared to other reactivation models used in the field. The stimulus of fur plucking is fast and easy and requires no specialized equipment. Further, dermatome skin lesions are easy to detect and measure and the model is applicable to both HSV-1 and HSV-2 and diverse mouse lines. Altogether the straightforward and tractable nature of this reactivation model makes it well-suited to investigating the pathogenic mechanisms of HSV, understanding the immune responses at play during recurrent disease, and evaluating vaccines and therapeutics. Furthermore, our observation that fur plucking the ipsilateral flank is sufficient to induce reactivation, whereas shaving or contralateral flank plucking did not induce reactivation, suggests that fur plucking triggers a response in innervating sensory neurons and provides an opportunity to study the neurobiology of HSV infection. Cell culture models of HSV latency in cultured neurons have defined stimuli such as axon damage, growth factor deprivation, and inflammatory cytokines as inducing the epigenetic changes required to activate viral gene expression from the latent genome (15–17). Future studies will investigate the effects of these and other stimuli in this new mouse model of HSV reactivation.

## ACKNOWLEDGEMENTS

This work was supported by R01 AI139512 and R01 AI175708 (H.M.L.). D.T.P. was supported by T32 AI007419 and F31 AI167502.

## SUPPLEMENTARY METHODS

### Viruses and cells

Virus stocks were grown in Vero (African green monkey kidney epithelial) cells. Titers of virus stocks were determined on Vero cells by a focus-forming assay (FFA). Vero cells were maintained in Dulbecco’s modified Eagle medium (DMEM) containing 5% heat-inactivated fetal bovine serum (FBS) and L-glutamine at 37°C with 5% CO_2_ containing 2% fetal bovine serum (FBS), L-glutamine, and HEPES at 37°C with 5% CO_2_. HSV-1 strain NS was obtained from Dr. Harvey Friedman (University of Pennsylvania) (18). HSV-2 strain 333 was obtained from Dr. Steven Bachenheimer (UNC). Virus stock titers were quantified by focus-forming assay on Vero cells. Viral foci were detected using 1:10,000 dilution of αHSV rabbit antibody (Dako #B0114) and 1:50,000 dilution of goat αrabbit HRP conjugated antibody (Sigma #12-348), and TrueBlue peroxidase substrate (KPL). Antibody incubations were performed for at least 1 hour at room temperature. Foci were counted on a CTL Immunospot analyzer.

### Mice

All experiments and husbandry were performed under the approval of the University of North Carolina at Chapel Hill Institutional Animal Care and Use Committee. Experiments used 8-12-week-old male and female mice on a C57BL/6 background, bred in-house. SKH-1 (Charles River strain #477) were received from Dr. Brian Conlon (UNC) and 10-12 week-old male and female mice were used for experiments.

### HSV skin infections

One day prior to infection, mice were anesthetized by nose-cone isoflurane and depilated by plucking on the right flank unless otherwise indicated. One day later, mice were anesthetized by chamber isoflurane for infections. To perform infections, we abraded the skin of anesthetized, depilated mice with ∼10 closely spaced punctures (∼5mm^2^ total area) using a Quintip allergy needle (Hollister Stier #8400ZA). Immediately after abrasion, we overlaid 10 µL of viral inoculum (virus + 1% FBS in PBS) and allowed the inoculum to dry while mice were anesthetized.

### Viral load measurements

Viral genomes were quantified from skin that was homogenized in 500 µL of PBS and silica beads on a MagNAlyser instrument (Roche). DNA was then extracted from 200 µL of homogenate using the Qiagen DNeasy Blood & Tissue Kit (#69504). Extracted HSV-1 genomes were then quantified by TaqMan qPCR on a CFX96 Touch real-time PCR detection system (Bio-Rad) against a standard curve generated by extracting DNA from an HSV-1 stock at 10^8^ FFU/mL and serially diluting. HSV-1 genomes were detected using the following primers against the UL23 gene: F primer 5′-TTGTCTCCTTCCGTGTTTCAGTT-3′, R primer 5′-GGCTCCATACCGACGATCTG-3′, and probe 5′-FAM-CCATCTCCCGGGCAAACGTGC-MGB-NFQ-3′ (19).

### Lesion area measurements

To measure HSV lesion areas, mice were anesthetized and photographed using an iPhone camera next to a ruler and an identifying card. Thereafter, images were analyzed using Image J in which pixels were converted to millimeters using the reference ruler and then lesions were outlined using the freehand tool and calculated areas within the freehand designations were reported.

## REFERENCES

1. Bradley H, Markowitz LE, Gibson T, McQuillan GM. 2014. Seroprevalence of herpes simplex virus types 1 and 2--United States, 1999-2010. J Infect Dis 209:325–333.

2. Kollias CM, Huneke RB, Wigdahl B, Jennings SR. 2015. Animal models of herpes simplex virus immunity and pathogenesis. J Neurovirol 21:8–23.

3. Suzich JB, Cliffe AR. 2018. Strength in Diversity: Understanding the Pathways of Herpes Simplex Virus Reactivation. Virology 522:81–91.

4. Shimeld C, Whiteland JL, Nicholls SM, Easty DL, Hill TJ. 1996. Immune cell infiltration in corneas of mice with recurrent herpes simplex virus disease. J Gen Virol 77 (Pt 5):977–985.

5. Shimeld C, Easty DL, Hill TJ. 1999. Reactivation of Herpes Simplex Virus Type 1 in the Mouse Trigeminal Ganglion: an In Vivo Study of Virus Antigen and Cytokines. J Virol 73:1767–1773.

6. Laycock KA, Lee SF, Brady RH, Pepose JS. 1991. Characterization of a murine model of recurrent herpes simplex viral keratitis induced by ultraviolet B radiation. Invest Ophthalmol Vis Sci 32:2741–2746.

7. Sawtell NM, Thompson RL. 1992. Rapid in vivo reactivation of herpes simplex virus in latently infected murine ganglionic neurons after transient hyperthermia. J Virol 66:2150–2156.

8. Yu W, Geng S, Suo Y, Wei X, Cai Q, Wu B, Zhou X, Shi Y, Wang B. 2018. Critical Role of Regulatory T Cells in the Latency and Stress-Induced Reactivation of HSV-1. Cell Rep 25:2379–2389.e3.

9. Vicetti Miguel RD, Sheridan BS, Harvey SAK, Schreiner RS, Hendricks RL, Cherpes TL. 2010. 17-β Estradiol Promotion of Herpes Simplex Virus Type 1 Reactivation Is Estrogen Receptor Dependent. J Virol 84:565–572.

10. Simmons A, Nash AA. 1984. Zosteriform spread of herpes simplex virus as a model of recrudescence and its use to investigate the role of immune cells in prevention of recurrent disease. J Virol 52:816–821.

11. van Lint A, Ayers M, Brooks AG, Coles RM, Heath WR, Carbone FR. 2004. Herpes simplex virus-specific CD8+ T cells can clear established lytic infections from skin and nerves and can partially limit the early spread of virus after cutaneous inoculation. J Immunol Baltim Md 1950 172:392–397.

12. Philip DT, Goins NM, Catanzaro NJ, Misumi I, Whitmire JK, Atkins HM, Lazear HM. 2023.Interferon lambda restricts herpes simplex virus skin disease by suppressing neutrophil-mediated pathology. BioRxiv Prepr Serv Biol 2023.09.11.557277.

13. Hill TJ, Blyth WA, Harbour DA. 1978. Trauma to the skin causes recurrence of herpes simplex in the mouse. J Gen Virol 39:21–28.

14. Benavides F, Oberyszyn TM, VanBuskirk AM, Reeve VE, Kusewitt DF. 2009. The hairless mouse in skin research. J Dermatol Sci 53:10–18.

15. Cuddy SR, Schinlever AR, Dochnal S, Seegren PV, Suzich J, Kundu P, Downs TK, Farah M, Desai BN, Boutell C, Cliffe AR. 2020. Neuronal hyperexcitability is a DLK-dependent trigger of herpes simplex virus reactivation that can be induced by IL-1. eLife 9:e58037.

16. Sawtell NM, Thompson RL. 2004. Comparison of herpes simplex virus reactivation in ganglia in vivo and in explants demonstrates quantitative and qualitative differences. J Virol 78:7784–7794.

17. Wilcox CL, Johnson EM. 1987. Nerve growth factor deprivation results in the reactivation of latent herpes simplex virus in vitro. J Virol 61:2311–2315.

18. Friedman HM, Macarak EJ, MacGregor RR, Wolfe J, Kefalides NA. 1981. Virus infection of endothelial cells. J Infect Dis 143:266–273.

19. Ma JZ, Russell TA, Spelman T, Carbone FR, Tscharke DC. 2014. Lytic Gene Expression Is Frequent in HSV-1 Latent Infection and Correlates with the Engagement of a Cell-Intrinsic Transcriptional Response. PLoS Pathog 10:e1004237.

